# Optimised biomolecular extraction for metagenomic analysis of microbial biofilms from high-mountain streams

**DOI:** 10.1101/2020.04.30.069724

**Authors:** Susheel Bhanu Busi, Paraskevi Pramateftaki, Jade Brandani, Stylianos Fodelianakis, Hannes Peter, Rashi Halder, Paul Wilmes, Tom J. Battin

## Abstract

Glacier-fed streams (GFS) are harsh ecosystems dominated by microbial life organized in benthic biofilms, yet the biodiversity and ecosystem functions provided by these communities remain under-appreciated. To better understand the microbial processes and communities contributing to GFS ecology, it is necessary to leverage high throughput sequencing. Low biomass and high inorganic particle load in GFS sediment samples may affect nucleic acid extraction efficiency using extraction methods tailored to other extreme environments such as deep-sea sediments. Here, we benchmarked the utility and efficacy of four extraction protocols, including an up-scaled phenol-chloroform protocol. We found that established protocols for comparable sample types consistently failed to yield sufficient high-quality DNA, delineating the extreme character of GFS. The methods differed in the success of downstream applications such as library preparation and sequencing. An adapted phenol-chloroform-based extraction method resulted in higher yields and better recovered the expected taxonomic profile and abundance of reconstructed genomes when compared to commercially-available methods. Affordable and straight-forward, this method consistently recapitulated the abundance and genomes of a “mock” community, including eukaryotes. Moreover, by increasing the amount of input sediment, the protocol is readily adjustable to the microbial load of the processed samples without compromising protocol efficiency. Our study provides a first systematic and extensive analysis of the different options for extraction of nucleic acids from glacier-fed streams for high-throughput sequencing applications, which may be applied to other extreme environments.

## Introduction

The advent of high-throughput sequencing technologies has brought hitherto inconceivable capacities to characterize the microbial ecology of both well-studied (Jansson and Hofmockel 2018; Nielsen and Ji 2015) and under-explored environments. Among the latter figure high-mountain and particularly glacier-fed streams (Milner et al. 2017) and the microbial biofilms that colonize their beds (Battin et al. 2016). Today, these streams are changing at an unprecedented pace owing to climate change and the thereby shrinking glaciers, and yet little is known of their microbial diversity (Wilhelm et al. 2013, Milner et al. 2017). Glacier-fed stream (GFS) sediments are extreme habitats characterized by low microbial cell abundance and activities but very high loads of fine mineral particles. In order to understand the diversity and composition of these microbial communities, including both eukaryotes and prokaryotes, and the role that they play, it is essential to extract nucleic acids in sufficient quantity and quality from often complex environmental matrices. After extracting the nucleic acids, downstream applications including molecular biology methods such as PCR and next-generation sequencing of amplicons or metagenomes allow for the compositional, functional and phylogenetic characterization of microbial populations and the communities that they form (Roume et al. 2013).

While there is no lack of protocols and literature pertaining to the extraction of nucleic acids from a wide variety of environments (Roume et al. 2013; Miller et al. 1999; Xin and Chen 2012; Porebski, Bailey, and Baum 1997; Zhou, Bruns, and Tiedje 1996), few reports dwell on the utility of these methods for biomolecular extractions from sedimentary samples with very low cell abundance as typical for GFS (Wilhelm et al. 2013; Ren, Gao, and Elser 2017). In 2015, Lever *et al.* elaborately described diverse factors and components that need to be considered for efficient nucleic acid extractions (Lever et al. 2015). These include, but are not limited to key steps like cell lysis, removal of impurities and inhibitors and of critical additives like carrier DNA molecules to enhance aggregation and thus precipitation of DNA in case of very low concentrations. Since the first extraction of DNA by Swiss medical doctor Friedrich Miescher in 1869 (Dahm 2008), biomolecule extractions have shifted from those performed with solutions prepared primarily in the laboratory (Sambrook and Russell 2006; Miller et al. 1999) to using commercially-available kits. These ready-made options are designed to avoid the use of volatile and toxic chemicals such as phenol and chloroform, and are tailored to various environments including blood, faecal material, plant and soils (Claassen et al. 2013; Psifidi et al. 2015; Smith, Diggle, and Clarke 2003; Vishnivetskaya et al. 2014). While studies have concentrated on nucleic acid extraction from glacial ice cores (Dancer, Shears, and Platt 1997) or surface snow (Pei-Ying et al. 2012), none has demonstrated their utility for GFS sediments. Together with low cell abundance, the complex mineral matrices in GFS - a consequence of the erosion activity of glaciers – may affect nucleic acid extraction efficiency. As we attempt to better understand how nature works at its limits through the study of extreme environments, non-commercial approaches and methodologies need to be revisited and optimized.

In recent years, several research groups (Besemer et al. 2012; Ren et al. 2017; Ren, Gao, and Elser 2017; Dancer, Shears, and Platt 1997) have successfully used kit-based methods for DNA extraction and subsequent 16S ribosomal RNA gene amplicon sequencing on GFS samples. However, the requirements for whole genome shotgun sequencing currently include at least 50 ng of input DNA to minimize bias due to PCR reactions during library preparation. Here, we address the utility and efficiency of the “gold” standard phenol-chloroform extraction (Dairawan and Shetty 2020), and three alternative methodologies to identify the process(es) that yield not only the highest quantity but also quality of DNA, from GFS sediments. We found that for the GFS sediment samples, the phenol-chloroform method yielded the expected diversity and taxonomic profiles, while also enabling reconstruction of metagenome-assembled genomes. Using a mock community, we additionally validated the utility of this method for the extraction of nucleic acids from both pro- and eu-karyotic sources. Overall, our findings provide a framework for the extraction of nucleic acids such as DNA for whole genome shotgun sequencing from GFS sediments, whilst highlighting the potential variability introduced due to the isolation method employed.

## Methods

### Sample origin & collection

DNA extraction protocols were benchmarked using three different GFS sediments from the Swiss Alps: Corbassière (CBS, 2444 m a.s.l) and Val Ferret (FE, 1995 m a.s.l) at the glacier snout (up site, FEU) and one kilometer further downstream (down site, FED). CBS differs from FEU and FED in terms of bedrock geology, with clastic sedimentary limestone dominating the catchment of CBS and Brecchia of gneiss dominating in FEU and FED. Sediments generally contain more organic material further downstream from the glacier, which may inhibit DNA extraction. Sediments (0.25 to 3.15 mm) were collected using flame-sterilized sampling equipment. Wet sediments were transferred into 10 ml sterile, DNA/DNase-free tubes and immediately flash-frozen in liquid nitrogen in the field. Samples were transferred to the laboratory and kept at −80 °C until analysis. All necessary measures were taken to ensure contamination-free sampling.

### DNA extraction methods

Four different DNA extraction methods were applied to the samples. The key characteristics of the different methods are summarized in Table 1. Method-1, −2 and −4 were manual protocols differing primarily in the lysis step (bead-beating and lysis buffer composition; (Roume et al. 2013; Lever et al. 2015; Sambrook and Russell 2006) while method-3 was a modified protocol of the DNeasy PowerMax Soil Kit (Cat.No. 12988-10) provided by Qiagen (based on communication exchanged with the manufacturer). Due to the very low microbial abundance additional precautions were taken to establish contamination-free conditions, including daily decontamination of equipment/areas with bleach, DNA/DNase-free glassware plasticware, reagents and chemicals.

**Table 1:**
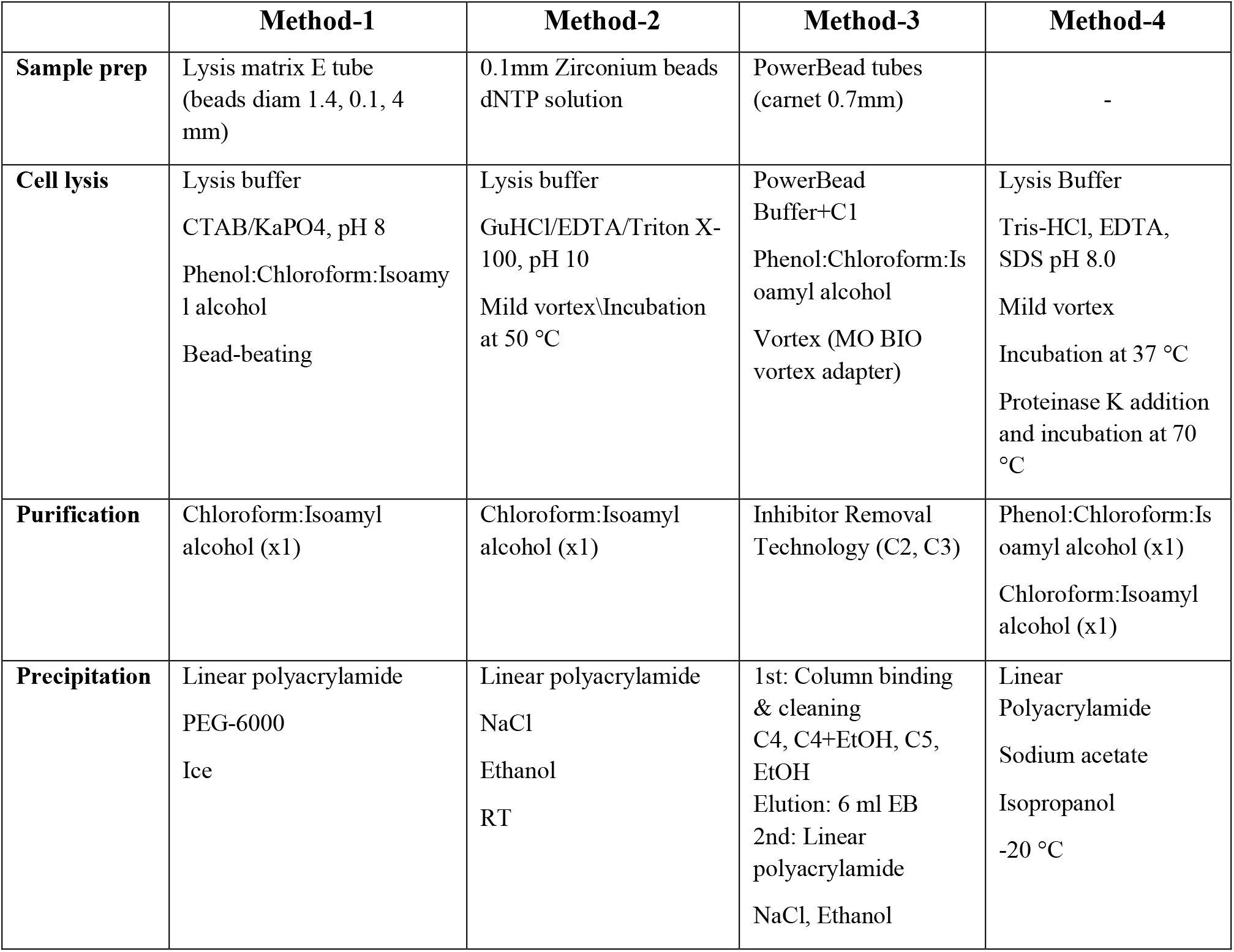
Key characteristics of the four selected methods

|  | Method-1 | Method-2 | Method-3 | Method-4 |
| --- | --- | --- | --- | --- |
| **Sample prep** | Lysis matrix E tube<br>(beads diam 1.4, 0.1, 4 mm) | 0.1mm Zirconium beads<br>dNTP solution | PowerBead tubes<br>(carnet 0.7mm) | - |
| **Cell lysis** | Lysis buffer<br>CTAB/KaPO<sub>4</sub>, pH 8<br>Phenol:Chloroform:Isoamyl alcohol<br>Bead-beating | Lysis buffer<br>GuHCl/EDTA/Triton X-100, pH 10<br>Mild vortex\Incubation at 50 °C | PowerBead Buffer+C1<br>Phenol:Chloroform:Is oamyl alcohol<br>Vortex (MO BIO vortex adapter) | Lysis Buffer<br>Tris-HCl, EDTA, SDS pH 8.0<br>Mild vortex<br>Incubation at 37 °C<br>Proteinase K addition and incubation at 70 °C |
| **Purification** | Chloroform:Isoamyl alcohol (x1) | Chloroform:Isoamyl alcohol (x1) | Inhibitor Removal Technology (C2, C3) | Phenol:Chloroform:Is oamyl alcohol (x1)<br>Chloroform:Isoamyl alcohol (x1) |
| **Precipitation** | Linear polyacrylamide<br>PEG-6000<br>Ice | Linear polyacrylamide<br>NaCl<br>Ethanol<br>RT | 1st: Column binding & cleaning<br>C4, C4+EtOH, C5, EtOH<br>Elution: 6 ml EB<br>2nd: Linear polyacrylamide<br>NaCl, Ethanol | Linear Polyacrylamide<br>Sodium acetate<br>Isopropanol<br>-20 °C |

Method-1 was based on a previously established method (Griffiths et al. 2000). Sample cell lysis was achieved by adding 0.5 g of sediment into a lysing matrix E tube with beads of variable diameter (SKU 116914050), 500 μl CTAB buffer (5% CTAB, 120 mM KPO_4_, pH 8.0) and 500 μl of phenol/chloroform/isoamyl alcohol (ratio 25:24:1). Samples were loaded on a Precellys beater for 45 s at 5.500 r/s. DNA was extracted once more with chloroform/isoamyl alcohol (24:1) and precipitated with 2 vol PEG-6000, 15 μg/ml linear polyacrylamide (LPA) and 2 h incubation on ice (Supplementary material).

Method-2 was an adaptation to alpine stream sediments of the modular method for DNA extraction previously published (Lever et al. 2015). Samples were prepared by mixing 5 g of sediment, 10-20% of 0.1mm zirconium beads and 1 ml of 100 mM dNTP solution. Cell lysis was achieved with 5 ml lysis buffer (30 mM Tris-HCl, 30 mM EDTA, 1% Triton X-100, 800 mM guanidium hydrochloride, pH 10.0) and incubation at 50 °C for 1h with gentle agitation. The supernatant was extracted once with chloroform/isoamyl alcohol (24:1) and DNA was precipitated with 10 μg/ml LPA, 0.2 vol 0.5 M NaCl, 2.5 vol ethanol and 2h incubation at RT in the dark (Supplementary material).

Method-3 has been previously applied successfully on sand and clay soils (Hale and Crowley 2015). In this protocol, the standard lysis capacity of the DNeasy PowerMax Soil Kit is supplemented and enhanced by the addition of phenol:chloroform:isoamyl alcohol along with PowerBeads (kit provided) and C1 solution. Then, an elaborate sequence of treatments and rinses with the standard buffers of the kit followed to reach elution of extracted DNA from silica columns with 6 ml of elution buffer. Further concentration of extracted DNA was carried out with the addition of 240 μl 5M NaCl, 2.5 vol ethanol and 10 μg/ml LPA (Supplementary material).

Method-4 involves chemical and enzymatic treatment of samples (Sambrook and Russell 2006). Five g of sample was mixed with 10 ml of lysis buffer (0.1 M Tris-HCl pH 7.5, 0.05 M EDTA pH 8, 1.25 % SDS) and 10 μl RNase A (100 mg/ml) with mild vortexing for 15 s and incubated at 37 °C for 1h in a hybridization oven. 100 μl Proteinase K (20 mg/ml) were added and incubation for 10 min at 70 °C. Samples were extracted once with phenol/chloroform/isoamyl alcohol (ratio 25:24:1) and supernatants were extracted once more with chloroform/isoamyl alcohol (24:1). DNA was precipitated with 10 μg/ml LPA, 1/10 vol 3M Sodium Acetate pH 5.2 and 1 vol isopropanol and overnight incubation at −20 °C (Supplementary material).

All DNA extracts were suspended in 100 μl of DNA/DNase-free water (ThermoFisher Scientific). Extracted DNA was thereafter stored at −20 °C until further use. Due to the low DNA yields, it was necessary to use a fluorescent DNA quantification method with high sensitivity like the Qubit dsDNA HS kit (Invitrogen). Quality assessment, with Nanodrop and DNA visualization on 0.8% agarose gel containing GelRed nucleic acid stain, was possible only for DNA extracted with method-4 and for DNA concentrations higher than 2 ng/μl. All samples yielded sufficient DNA for metagenomic sequencing and subsequent analyses.

### DNA sequencing

All DNA samples were subjected to random shotgun sequencing. The sequencing libraries were prepared using the NEBNext Ultra II FS DNA Library Prep kit for Illumina (Cat.No. E7805) using the protocol provided with the kit. The libraries were prepared considering 350 basepairs (bp) average insert size. Qubit (Invitrogen) was used to quantify the prepared libraries while their quality was assessed on a Bioanalyzer (Agilent). We used the NextSeq500 (Illumina) instrument to perform the sequencing using 2×150 bp read length at the Luxembourg Centre for Systems Biomedicine Sequencing Platform.

### Genome reconstruction and metagenomic data processing

Paired sequences (i.e., forward and reverse) were processed using the Integrated Meta-omic Pipeline (IMP) (Narayanasamy et al. 12/2016). The metagenomic workflow encompasses pre-processing, assembly, and genome reconstruction in a reproducible manner. The adapter sequences were trimmed in the pre-processing step including the removal of human reads. Thereafter, *de novo* assembly was performed using the MEGAHIT (version 2.0) assembler^(D. Li et al. 2015)^. Default IMP parameters were retained for all samples. Subsequently, we used MetaBAT2 (Kang et al. 2019) and MaxBin2 (Wu, Simmons, and Singer 2016) for binning in addition to an in-house binning methodology previously described (Heintz-Buschart et al. 2017). The latter method initially ignores the ribosomal RNA sequences in kmer profiles based on VizBin embedding clusters (Laczny et al. 2015). In this context, VizBin utilises density-based non-hierarchical clustering algorithms and depth of coverage for genome reconstructions. Subsequently we obtained a non-redundant set of metagenome-assembled genomes (MAGs) using DASTool (Sieber et al. 2018) with a score threshold of 0.7 for downstream analyses. The abundance of MAGs in each sample was determined by mapping the reads to the reconstructed genomes using BWA-MEM (H. Li 2013), taking the average coverage across all contigs. Diversity measures from metagenomic sequencing were assessed by determining the abundance-weighted average coverage of all the reads to identify the number of non-redundant read sets (Rodriguez-R and Konstantinidis 2014).

### Taxonomic classification for metagenomic operational taxonomic units

We used the trimmed and pre-processed reads from the IMP workflow to determine the microbial abundance and taxonomic profiles based on the mOTU (v2) tool (Milanese et al. 2019). Based on the updated marker genes in the mOTU2 database including those from the TARA Oceans Study (Sunagawa et al. 2015) and recently generated MAGs (Tully, Graham, and Heidelberg 2018), taxonomic profiling was performed on our sequence datasets. We used a minimum alignment length of 140 bp to determine the relative abundances of the mOTUs, including the normalisation of read counts to the gene length, also accounting for the base coverage of the genes. Additionally, we used CheckM (Parks et al. 2015) to assess completeness and contamination. Subsequently, taxonomy for MAGs recovered after the redundancy analyses from DASTool was determined using the GTDB (Genome Taxonomy Database) toolkit (gtdb-tk) (Parks et al. 2018).

### Data analysis

All figures for the DNA concentrations, library preparation, assembly metrics and supplementary figures were generated using GraphPad Prism (v8.3.0). Taxonomical assessment and diversity measures were created using version 3.6 of the R statistical software package (Team 2013). DESeq2 (Love, Huber, and Anders 2014) with FDR-adjustments for multiple testing were used to assess significant differences in the MAG abundances. The genomic cluster figure for the mock community was obtained as an output from the IMP metagenomic pipeline.

## Results

### Phenol-chloroform-based extraction method results in higher DNA yields

To ensure native sequencing, by minimizing the number of PCR (polymerase chain reaction) cycles within the library preparation protocols, we tested four protocols for biomolecular extraction, with an aim of acquiring large quantities (>50 ng) DNA from glacier-fed stream benthic sediments. The four methods tested were selected because of their wide applicability on related environmental samples (Method-1 & −2) (Griffiths et al. 2000; Lever et al. 2015; Tatti et al. 2016) and their improved chances of higher yields (Method-3; Qiagen communication). Since method-4 is considered the gold standard of DNA extraction in biomedical sciences (Dairawan and Shetty 2020) and of bacterial cultures (Green and Sambrook 2017), it was included in our study. The four protocols are largely based on the same principles, *viz*. sample preparation, cell lysis, purification, precipitation and washing (Table 1). From preliminary tests, it became apparent that a small-scale approach (0.5 g input sediment) did not yield sufficient amounts of DNA for metagenomics due to, on average, limited microbial biomass in the samples. Thus, all protocols (aside from Qiagen’s - already produced for maxi scale) were scaled up to 5 g of input sediment and a co-precipitant, like linear polyacrylamide, was included in all precipitation steps. This was essential for the quantitative recovery of the small amounts of extracted DNA from high solution volumes (6-10 ml).

Overall, we found that extractions using the commercial kit from Qiagen (method-3) yielded increased total DNA as compared to a commonly used protocol (method-1; Fig. 1A). Furthermore, method-3 was similar in terms of DNA yield when compared to a generalized protocol (method-2) previously proposed (Lever et al. 2015) (Fig. 1B). On the other hand, the phenol-chloroform based extraction protocol (method-4) was tested against both methods 2 and 3, using sediment samples collected from the three different glacier floodplain streams (CBS, FEU, FED) from Switzerland. Method-1 was omitted from these tests due to insufficient DNA concentrations in the preliminary extractions. We found that for all three GFS, the phenol-chloroform extraction yielded the highest DNA concentrations. In some cases, and notably samples with low cell abundance, we even obtained one order of magnitude more DNA (Fig. 1C).

**Figure 1.**
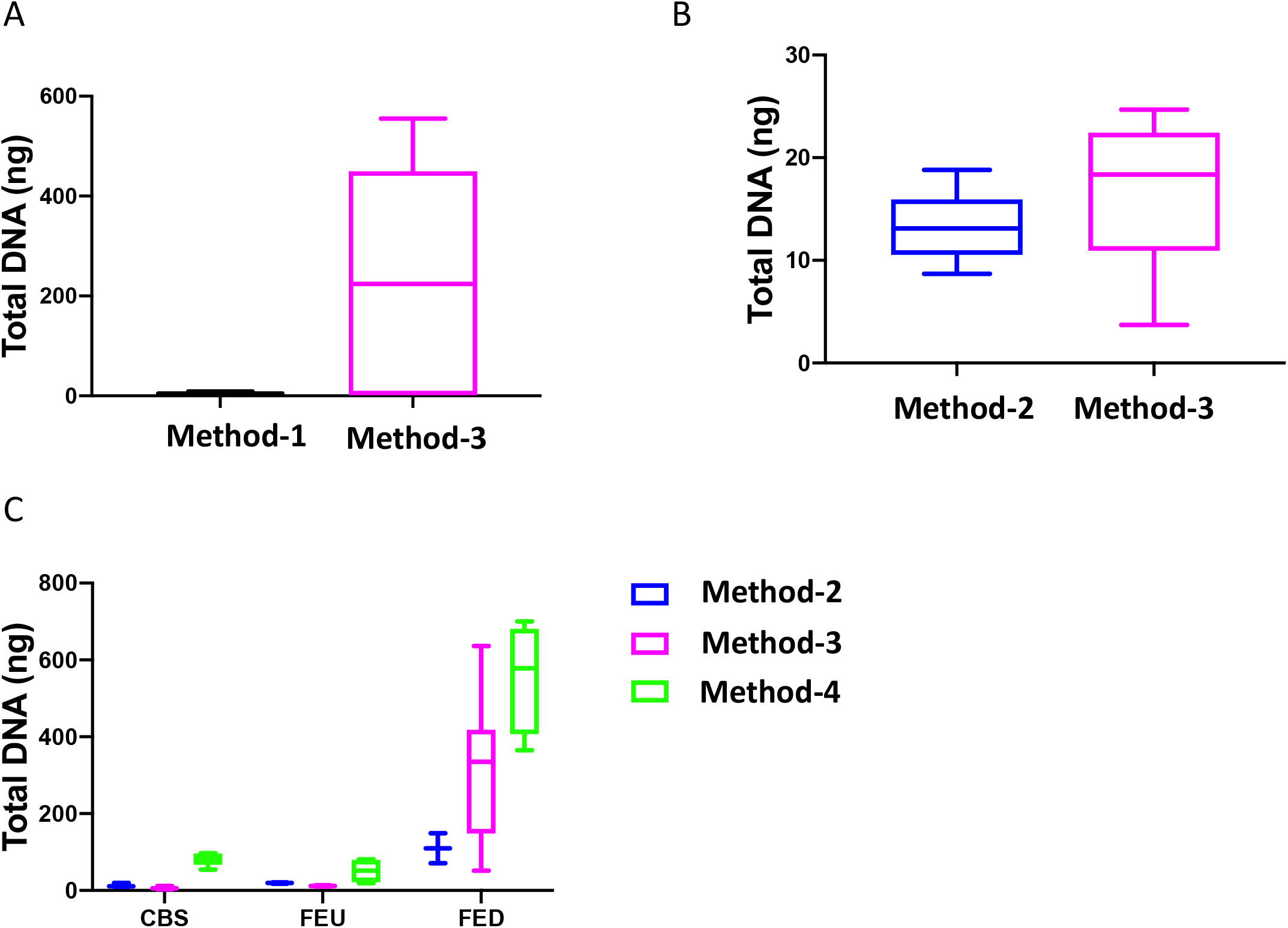
Total DNA concentrations using different extraction protocols. Boxplots represent the total amount of DNA (ng) extracted from 5 g of sediment when comparing (A) method-1 versus the modified-commercial kit-based method-3 and (B) method-2 versus method-3. (C) Boxplots of the DNA quantities isolated from three glacial floodplains (CBS - Corbassière, FEU - Val Ferret up site, FED - Val Ferret down site), using method −2, −3 and −4.

Quality assessment of these DNA extracts with Nanodrop showed OD260/280 ratios in between ~1.4 and ~1.6. Agarose gel electrophoresis revealed a high-molecular weight band with no apparent shearing, smearing or residual RNA, indicative of high-quality DNA (Fig. 2). A secondary effect appearing in certain samples, but without any perceived consequences in the quality of extracted DNA whatsoever, was the development of a pink-fuchsia color of varying intensities with the addition of phenol:chloroform:isoamyl alcohol (Fig. 3). This was pH dependent since samples were decolorized with the addition of sodium acetate pH 5.2 in the precipitation step. This could possibly be due to a ferric-chloride-phenol compound formed when chloride and phenol constituents of the protocol interact reversibly with Fe^+3^ ions contained in certain samples depending on local geology (Banerjee and Haldar 1950). Similar coloration has been previously reported (Lever et al. 2015).

**Figure 2.**
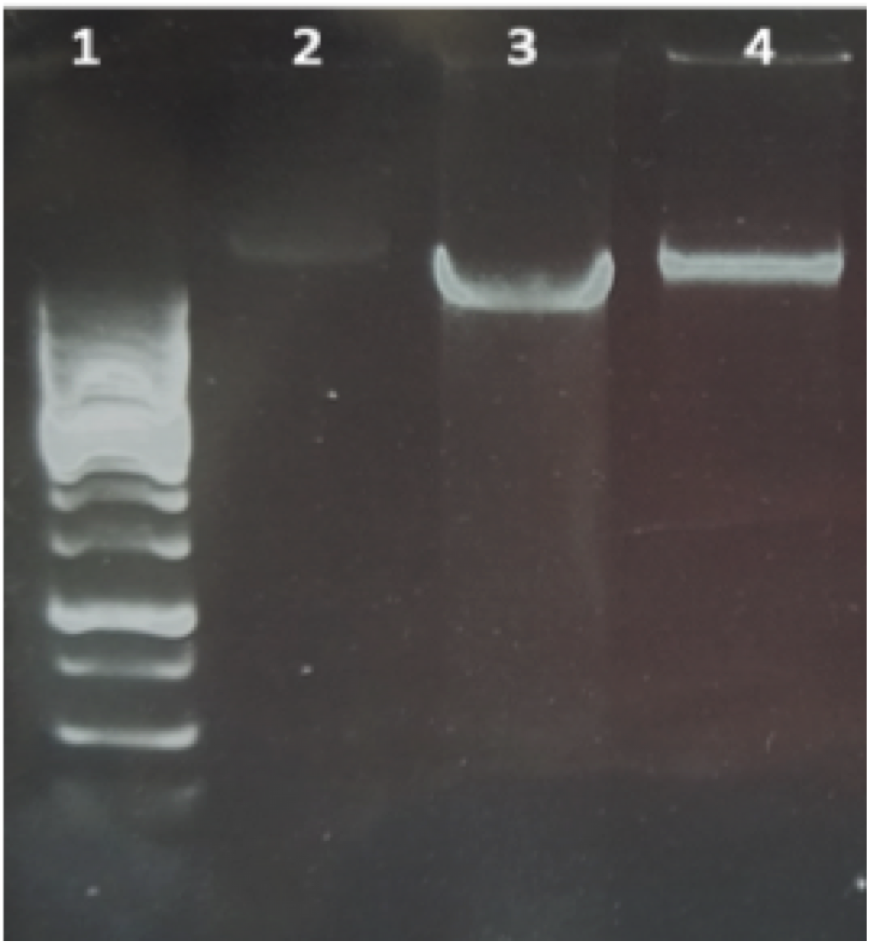
Agarose gel electrophoresis of DNA extracted with method-4. Lane 1: GeneRuler 1 kb DNA ladder; lanes 2-4: CBS, FED, FEU respectively.

**Figure 3.**
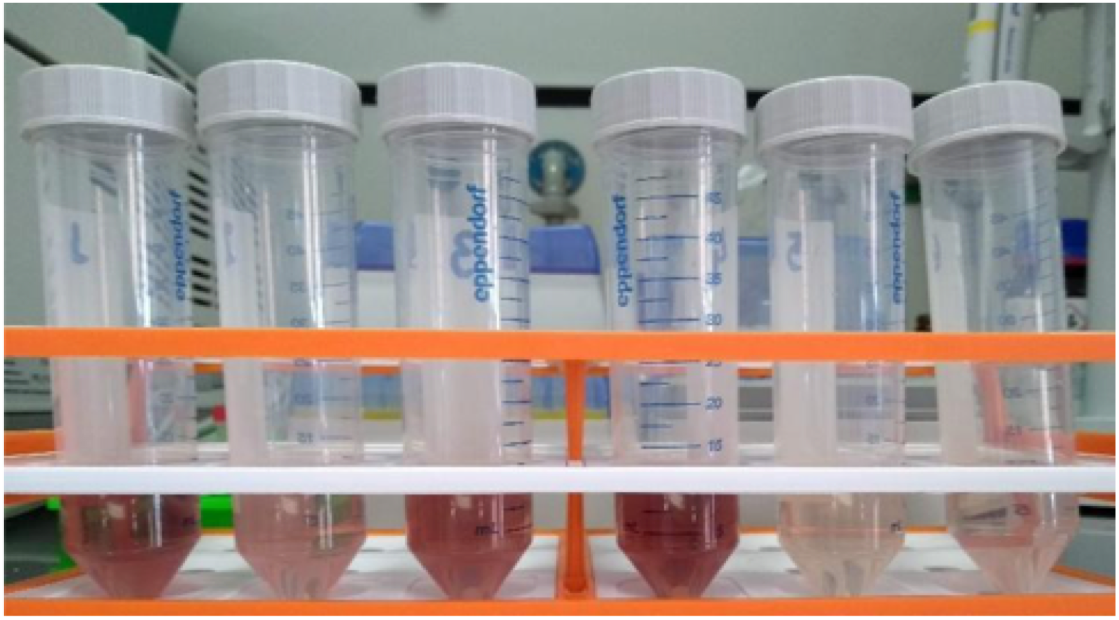
Pink-fuchsia supernatants developed during phenol:chloroform extraction step

### Extraction method affects library preparation efficiency

The DNA extracted based on method-3 and using phenol-chloroform methods were subsequently subjected to library preparation for high-throughput whole genome shotgun sequencing. Despite similar quality of DNA across both methods (~1.4-1.6 OD_260/280_), library preparation using the modified commercial kit did not yield any successful libraries (Fig. 4). To assess if any impurities or inhibitors hampered library preparation we tested two clean-up methods for the DNA extracted from the commercial kit: 1) ethanol precipitation and 2) magnetic-bead based clean-up. We found that the magnetic-bead method leads to a complete loss of sample (i.e., undetectable DNA quantity via Qubit analyses) during the process, especially if starting with a low input DNA concentration. Although we lost six out of twelve samples using the magnetic-bead clean-up, we achieved 100% efficiency in library preparation with the remaining six samples. On the other hand, ~20% of the samples cleaned via ethanol precipitation failed library preparation. Contradictory to these methods, DNA extracted using the phenol-chloroform based method (method-4) yielded 100% efficiency in terms of library preparation without any additional clean-up (Fig. 4). Additionally, we found that the distribution of the total yield after library preparation using the phenol-chloroform method was more uniform across samples compared to the other methods (Fig. 4).

**Figure 4.**
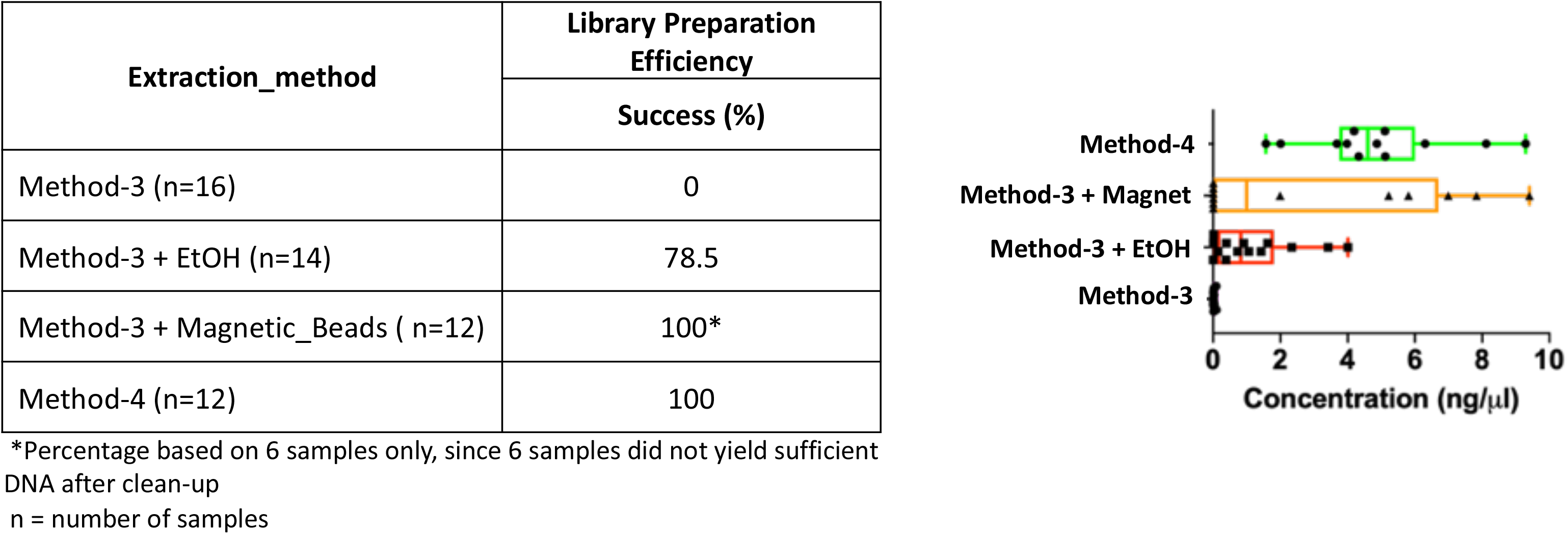
Library preparation efficiency. The efficiency or success percentage for prepared libraries based on the individual methods is indicated in the table. Boxplots represent concentrations of the prepared libraries.

### Whole genome shotgun assembly unaffected by extraction methods

Extraction methods for whole genome shotgun sequencing may affect the sequencing itself, including the quality and assembly of the reads downstream. To assess this, we used the libraries prepared as described above (Fig. 4), and performed whole genome shotgun sequencing on an Illumina NextSeq500. The average quality across all three methods based on short-read sequencing was Q30 after trimming the leading and trailing sequences (described in Methods). We assessed several assembly metrics including the average length of contigs (N50), largest alignment, total aligned length and coverage. We did not find any significant differences among any of these measures across all three methods (Fig. 5A-C, 5E). Using a diversity index metric, we however found a more uniform distribution across all samples prepared using method-4, albeit no significant differences were observed compared to the commercial kit-based extraction and library preparation (Fig. 5D).

**Figure 5.**
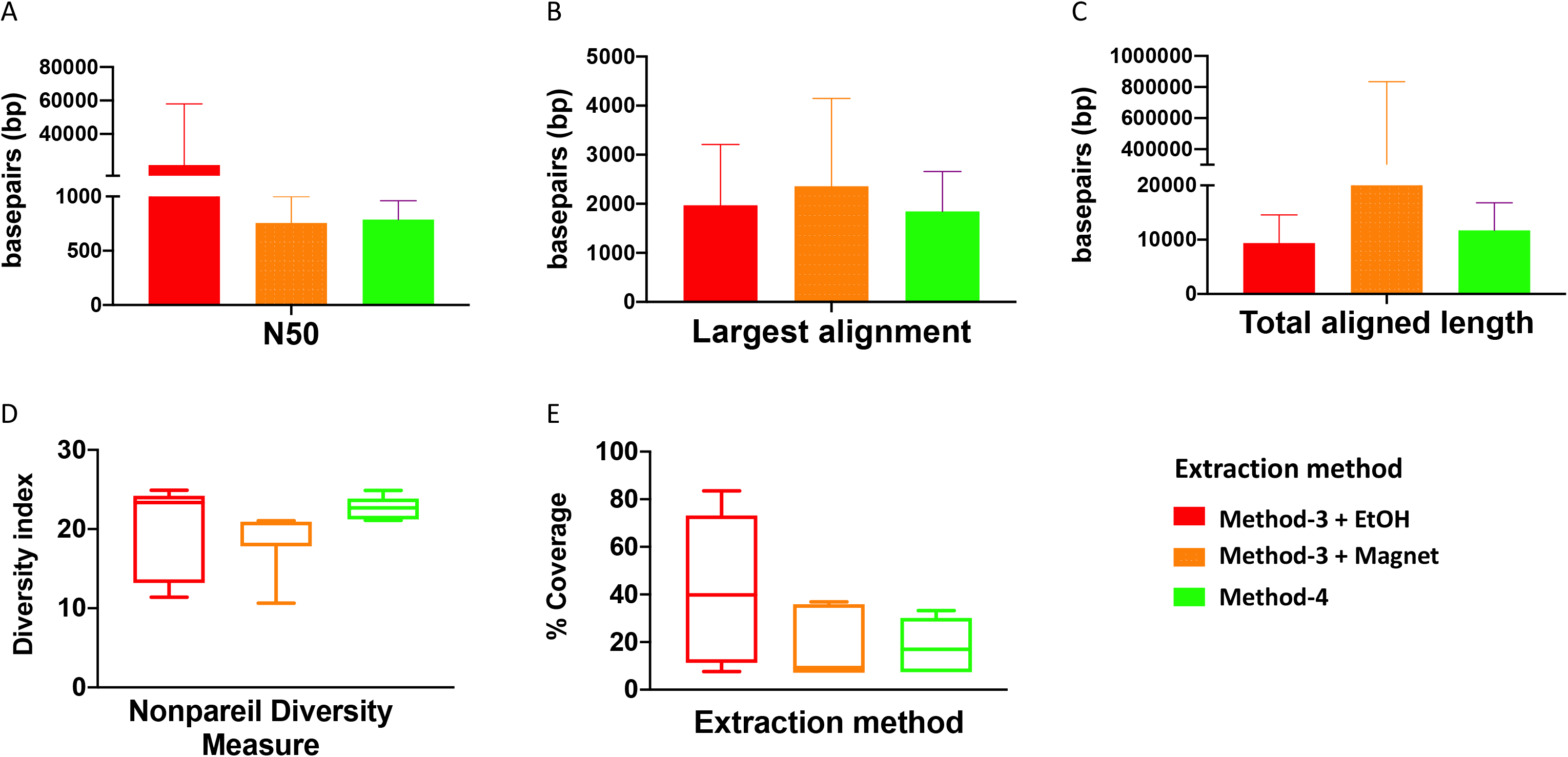
Estimate of assembly metrics following extraction. Barplots demonstrate the (A) N50 for the sequence assemblies, (B) length of the longest aligned sequence, (C) the total aligned length. Bars indicate standard deviation from the mean. (D) Boxplot showing the nonpareil diversity index across the three groups. (E) Percentage of coverage of the assembled sequences by read-mapping is depicted.

### Extraction methods influence metagenomic profiles

It is well established that extraction methods (Wagner Mackenzie, Waite, and Taylor 2015) and library preparation (Bowers et al. 2015) protocols affect the taxonomic profiles and genomes recovered after high-throughput sequencing. We determined if the preparation methods affected the overall diversity of taxa recovered and found that phenol-chloroform and the magnetic-bead clean-up methods demonstrated similar levels of diversity (Shannon) as compared to samples precipitated using ethanol (Fig. 6A). Overall, the community profiles of the ethanol precipitation-based method were highly diverse (Fig. 6B). Interestingly, the genomes recovered and their abundances were similar in the phenol-chloroform and magnetic-bead methods as well (Fig. 6C). However, we observed a significant increase (*p*<0.001, FDR-adjusted *p*-value) in the abundance of a *Ralstonia* genome when prepared with the ethanol precipitation protocol (Supplementary fig. 1). Additionally, we found that the number of genomes recovered using the phenol-chloroform was more consistent with previously reported 16S rRNA gene sequencing profiles for GFS from Austria (Wilhelm et al. 2013; Besemer et al. 2012; Wilhelm et al. 2014). Simultaneously, we used an approach to identify metagenomic operational taxonomic units (mOTUs) and found that the phenol-chloroform and magnetic-bead methods showed similar profiles of mOTUs compared to that of ethanol precipitation (Fig. 6D).

**Figure 6.**
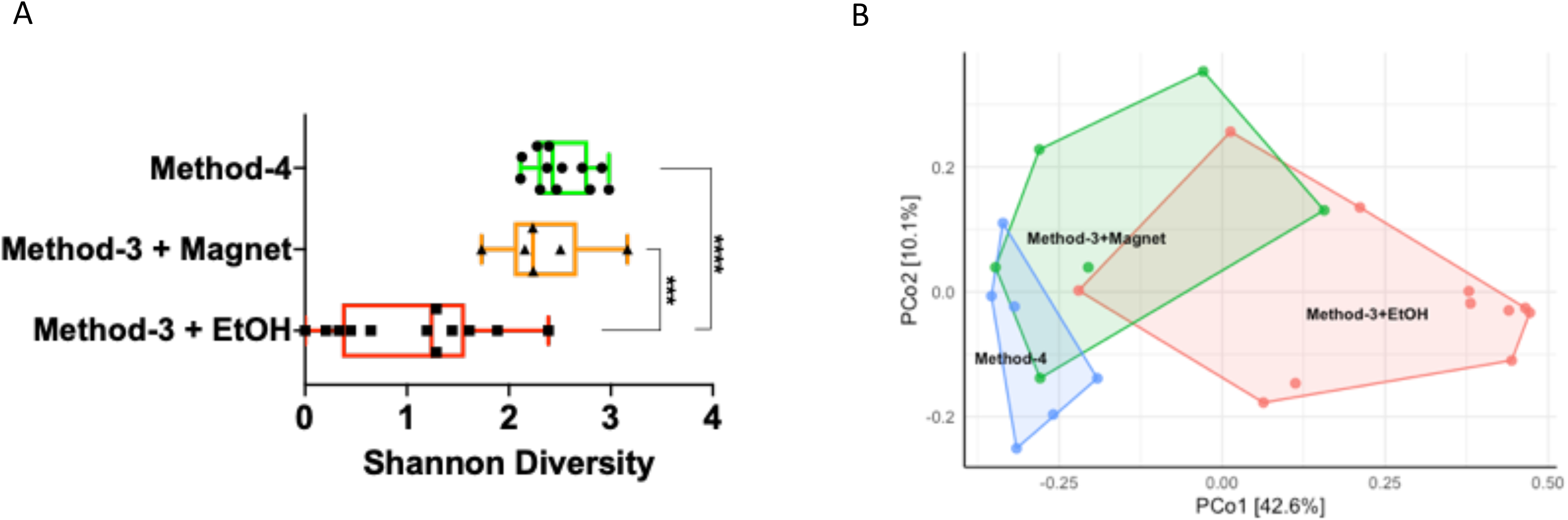
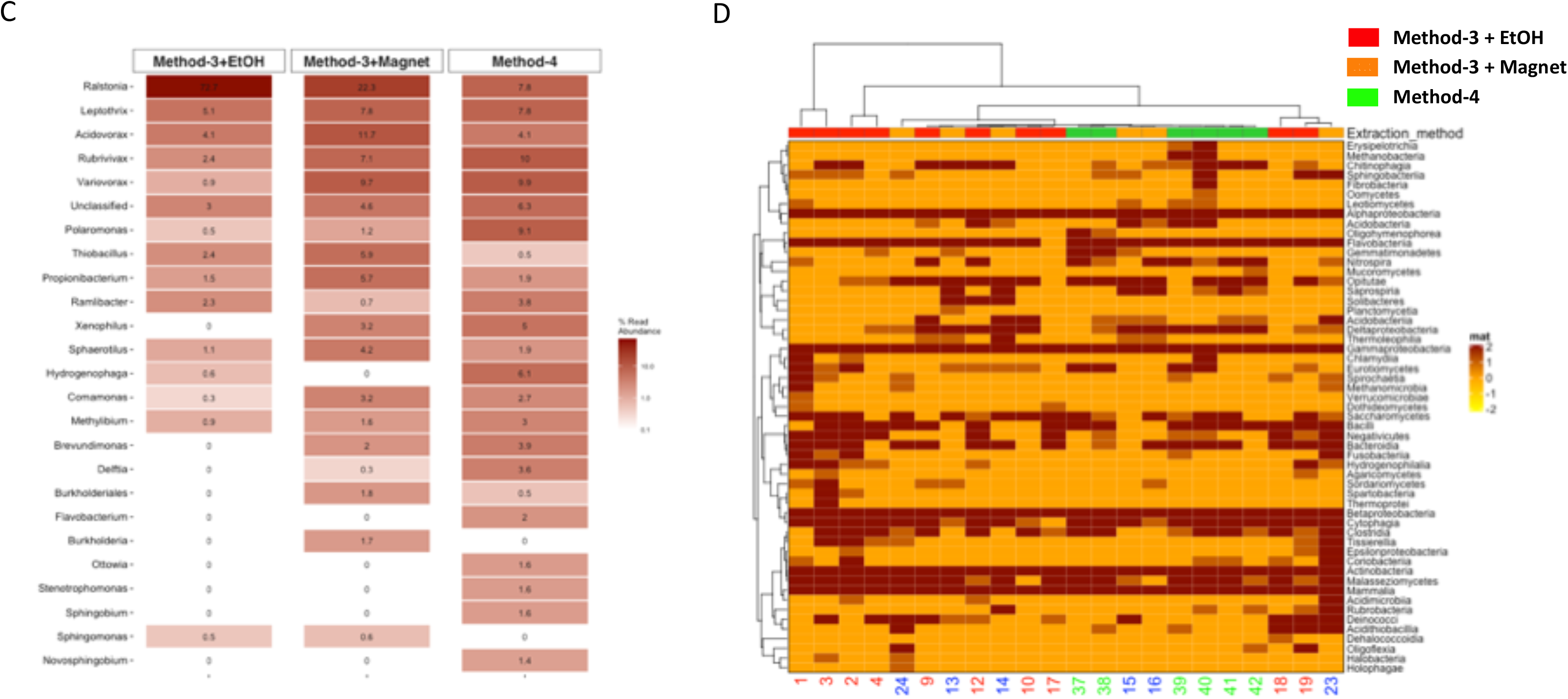
Diversity and taxonomic profiles of the metagenomic sequencing. (A) Boxplot showing the Shannon diversity index for the taxonomic profiling for the three groups. Significance was tested using a One-way ANOVA with Student-Neuman Keul’s post-hoc analysis. \*\*\**p*-value<0.001, \*\*\*\**p*-value<0.0001. (B) Principal component analyses generated using Bray-Curtis dissimilarity matrix depicts similarities or lack thereof between the three groups. (C) Abundances of the reconstructed genomes are depicted for method-3 + EtOH, method-3 + magnetic bead clean-up and method-4 extraction. (D) Heatmap demonstrating the mOTUs for the three methods is depicted. The hierarchical clustering for the heatmap was generated using Ward’s clustering algorithm.

### Efficiency of phenol-chloroform extraction on a mock community including eukaryotes

To determine whether the phenol-chloroform extraction method is biased against eukaryotes, we used a commercially-available mock community (ZymoBIOMICS Microbial Community Standard #D6300) to assess bias and errors. After sequencing, we recovered high quality (>90% completion, <5% contamination) bacterial genomes (Fig. 7A). Additionally, the abundance of the microbial genomes, including one of the eukaryotes - *Saccharomyces cerevisiae*, were similar to the expected levels in the mock community (Fig. 7B). On the other hand, the protocol enabled the identification and partial recovery of the *Cryptococcus neoformans* genome, albeit at lower levels possibly due to increased melanisation of the cell wall (Grossman and Casadevall 2017) affecting lysis and subsequent extraction.

**Figure 7.**
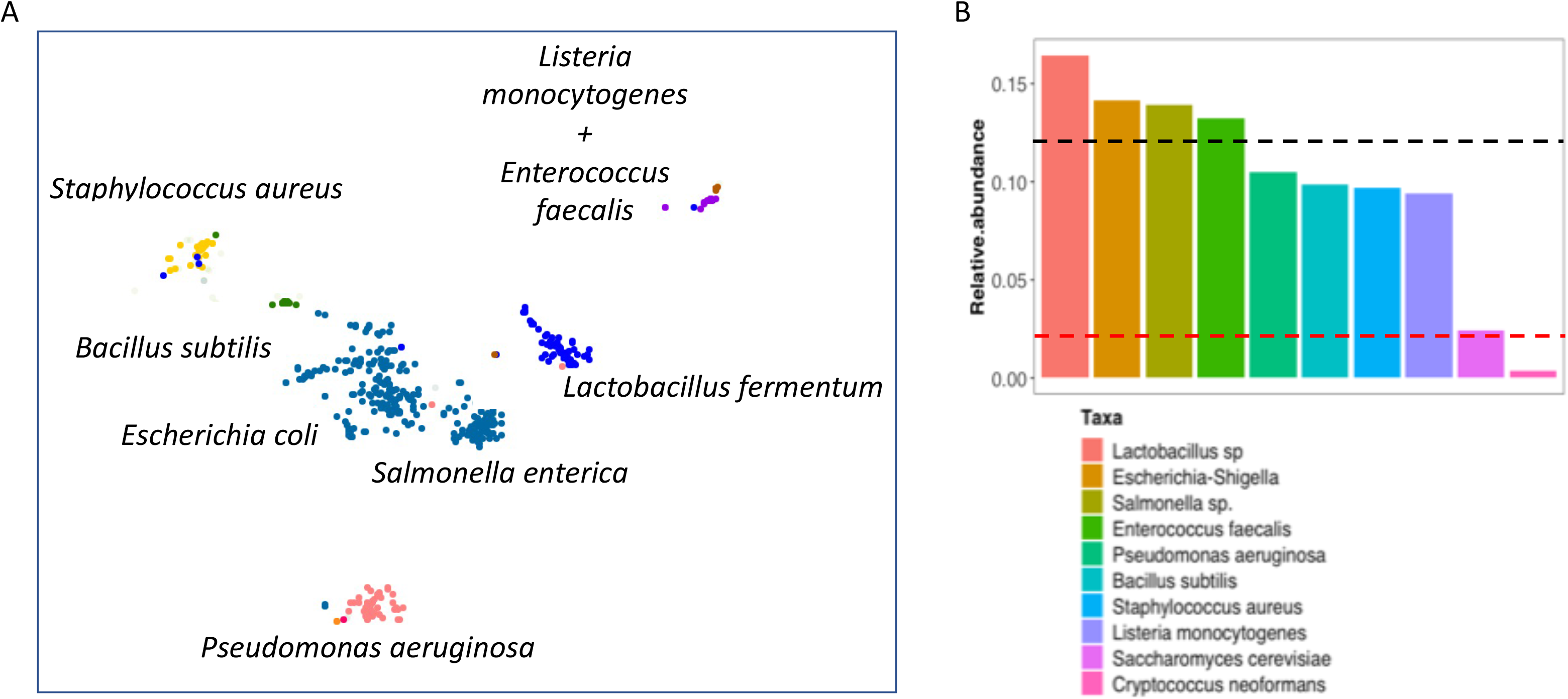
Evaluation of phenol-chloroform extraction using a mock community. (A) Scatterplot depicts the clusters of contigs representative of the reconstructed genomes after processing the mock community using the IMP meta-omics pipeline. The taxonomic identity is displayed next to the respective clusters. (B) Barplots indicate the relative abundance of the individual genomes recovered from the mock community sequencing after extraction with the phenol-chloroform method. The upper (black) line represents the expected abundance (12%) of the prokaryotes, while the lower (red) line indicates the expected abundance (2%) of the eukaryotes.

## Discussion

Improved omic techniques not limited to metagenomics are robust methods for analyzing nucleic acids and the characterisation of microbial communities in various environments (Jansson et al. 2012). One way of understanding the impacts of global climate change on GFS includes the establishment of their census of microbial life (Milner et al. 2017). However, methods designed for the extraction of biomolecules including DNA have not been validated for GFS sediments. Although previous glacier-fed streams studies successfully used extracted DNA for 16S rRNA amplicon sequencing (Ren et al. 2017; Ren, Gao, and Elser 2017; Vardhan Reddy et al. 2009; Wilhelm et al. 2013) the input DNA concentration requirements are considerably higher for whole genome shotgun sequencing. In order to pursue a deeper characterisation of the microbial communities within the GFS sediments, increased concentrations of DNA may additionally alleviate PCR biases (Brooks et al. 2015; Kim and Bae 2011). Also, as previously highlighted, several methods exist for extractions from a wide variety of environmental samples, but not for GFS sediments. Here, we systematically tested the utility of four extraction protocols to identify a ubiquitous methodology. We found that a phenol-chloroform based extraction protocol can be used for samples across geographical separations, differences in bedrock, and samples collected at various distances from the glacier.

Glassing *et al*. demonstrated that inherent DNA contamination may influence microbiota interpretation in low biomass samples (Glassing et al. 2016). Additionally, it is known that certain compounds - polysaccharides, humic acids, may affect PCR reactions (Rådström et al. 2004), requiring the need for additional DNA clean-up. It has been established that DNA losses occur during the purification steps (Roose-Amsaleg, Garnier-Sillam, and Harry 2001), including when using commercial column methods (Howeler, Ghiorse, and Walker 2003; Lloyd et al. 2013), and phenol-chloroform (Ogram, Sayler, and Barkay 1987). Interestingly, we found similar losses when using the magnetic bead clean-up, whereas the ethanol precipitation method was inefficient compared to the phenol-chloroform protocol. Though the kit-based methods are more convenient and safer than phenol-chloroform extractions (Tesena et al. 2017), access to reagents and costs may be a considerable factor. On the other hand, isolation of the aqueous phase from phenol-chloroform can be user-dependent potentially affecting reproducibility, while kits have been shown to be more consistent across samples (Claassen et al. 2013). Another key feature of our findings was the potential for the kit-based methods to influence the efficiency of genome reconstruction and variability in the taxonomic profiles that were recovered. While this has been reported previously (Wagner Mackenzie, Waite, and Taylor 2015; Carrigg et al. 2007), we found considerable variability when compared to the phenol-chloroform. This is plausible due to the incomplete dissolution of DNA in buffers, especially when using methods involving charged minerals (Vorhies and Gaines 2009; Barton et al. 2006; Vishnivetskaya et al. 2014), which may additionally affect DNA stability. Additionally, we found that method-4 may be used for low biomass sediment samples and can be scaled down in the context of abundant cells per gram (data not shown).

The utility of extraction methods extends beyond the process itself, impacting downstream applications such as whole genome shotgun sequencing. Our study shows that phenol-chloroform may be an under-appreciated yet powerful method for isolating nucleic acids from glacier-fed stream sediments. While additional steps may be required towards extraction of other biomolecules such as RNA, proteins and metabolites, minor modifications may be sufficient (Toni et al. 2018). Moreover, we report for the first time a systematic assessment of biomolecular extraction methods for GFS sediments. Our findings though fundamental and previously unexplored, may lay the foundations for future in-depth characterisation of GFS microbial communities.

## Data Availability

The sequencing data generated during the current study are available from NCBI under the BioProject accession number PRJNA624048. A reporting summary for the uploaded data has been included as a metadata file at the accession listed ID. All extraction protocols including the modified commercial methods are available in the *Supplementary Materials*.

## Acknowledgments

We are grateful to Laura de Nies, Camille-Martin Gallausiaux, Jean-Pierre Trezzi, Cedric Laczny, Audrey Frachet, Lea Grandmougin, Annegrat Daujemont, Laura Lebrun (LCSB) for discussions and laboratory support. The University of Luxembourg Sequencing Platform and HPC facilities were highly instrumental for the *in-silico* analyses. The present work was supported by NOMIS Foundation and the Swiss National Science Foundation (CRSII5_180241) to Tom J. Battin.

## Contributions

P.P. performed the biomolecular extractions, including the validation of methods 1-3 alongside quality analyses and quantification. S.B.B. curated and validated the phenol-chloroform extraction method and whole genome shotgun sequencing analyses. T.B. and H.P. collected the glacier-fed stream samples for the experiments. S.F. did the DNA extractions, quantification and qualifications alongside P.P. R.H. handled the library preparation for all samples and the subsequent sequencing. P.W. contributed significantly to the development of method-1 in the manuscript. S.B.B., P.P., S.F., H.P., P.W., and T.J.B. conceived and formulated the experiments. S.B.B. and P.P. developed the manuscript with equal contributions from all authors.

## Ethical declarations

## Conflicts of interest

The authors do not have any competing interests.

---

**Supplementary figure 1.**
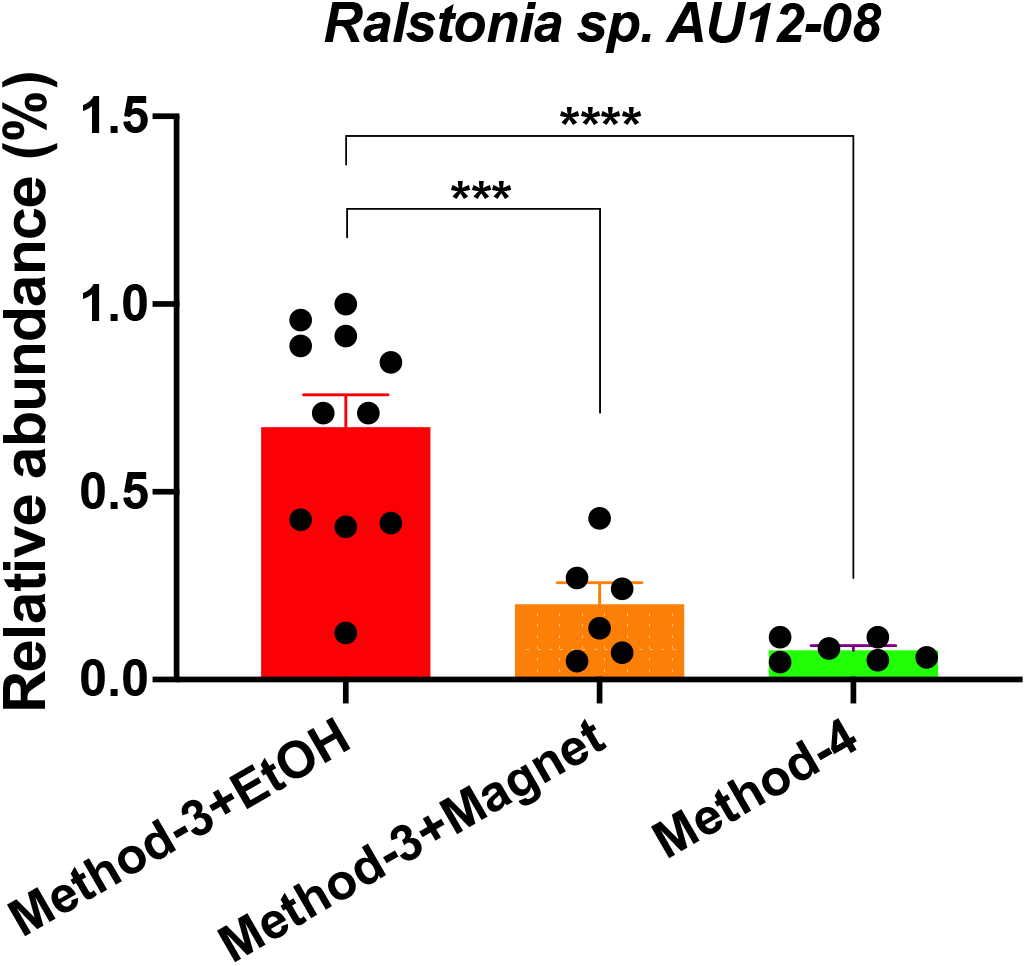
Relative abundance of Ralstonia sp. AU12-08. The abundance of the *Ralstonia* genome recovered from the samples when processed with method-3 (EtOH and magnetic bead clean-up) and method-4. Significance was tested using One-Way ANOVA with Student-Neuman Keul’s post-hoc analyses. ***p<0.001, ****p<0.0001

## Notes

https://dataview.ncbi.nlm.nih.gov/object/PRJNA624048?reviewer=oeqone2lcnbjcosvas3a9k3nu3

### Competing Interest Statement

The authors have declared no competing interest.

