## Supplementary Information for "Optimised biomolecular extraction for metagenomic analysis of microbial biofilms from high-mountain streams"

### Optimising biomolecular component extraction for meta-omic sequencing of microbial biofilms from high-mountain streams

#### Extraction protocols

##### Method -1

1. Add 0.4 g of the sample in a lysis matrix E tube, 500 µl CTAB Extraction buffer (5% CTAB, 120 mM $KPO_4$, pH 8) and bead beat in the Precellys at 5.500 r/s for 5 sec.
2. Add 500 µl of Phenol:Choloform:Isoamyl alcohol (25:24:1) close the tubes very carefully and bead beat them in the Precellys at 5.500 r/s for 45 sec.
3. Centrifuge for 10 min at 13000 g, at 4 °C.
4. Transfer the supernatant (app. 600 µl) to a 2-ml tube and extract with 1vol of chloroform:Isoamyl alcohol (24:1).
5. Centrifuge for 5 min at 13.000xg, at 4 °C.
6. Transfer the supernatant (600 µl) to a 2-ml tube.
7. Add 5.4 µl of linear acrylamide (15 µg/ml in 1800 µl total) and extract with 2 vol of PEG-6000 and precipitate for 2 h on ice.
8. Centrifuge for 60 min at 13.000xg, at 4°C.
9. Decant and remove rest carefully with a pipette.
10. Add 1000 µl of ice-cold 70% Ethanol and vortex briefly.
11. Centrifuge for 10 min at 13.000xg, at 4 °C.
12. Decant and remove rest carefully with a pipette.
13. Wash with 70% ethanol once more.
14. Dry the total nucleic acid for 5 min at RT (or longer until residues of alcohol are evaporated)
15. Elute pellet in 50 µl of molecular grade water

##### Method-2

*IMPORTANT: Keep sediment cold throughout the extraction procedure.*

1. Add up to 5 g of sediment per 15 ml bead-beating tube filled to 10-20% with 0.1-mm Zirconium beads per volume.
2. Add 1 ml of 10 mM dNTP solution. Gently shake to entirely soak or coat the sample.

*Purpose: dNTP has similar sorption characteristics to DNA. Coating sediment/mineral surfaces with dNTP thus reduces DNA sorption and drastically enhances DNA recovery from mineral-rich, oligotrophic samples*

3. Add 5 mL lysis solution I (30 mM Tris HCl, 30 mM EDTA, 800 mM guanidine hydrochloride, 0.5 % Triton X-100, final pH 10), and homogenize by inverting and tapping. Vortex briefly.
4. Place tubes in horizontal holder on Vortex Genie and shake at maximum speed for 30s. Incubate at 50 °C for 1 hour in a hybridization oven with rotation.
5. Spin down sediment at 10,000×g and 4 °C for 10 min and transfer supernatant to clean Eppendorf tube. Transfer as much supernatant as possible.
6. Add 1 volume of cold 24:1 chloroform-isoamyl alcohol mixture to supernatant. Mix by vortexing for 10 s and spin at 4 °C for 10 min at 10,000×g. Repeat this wash once, taking care to avoid any chloroform and particle transfer after second wash.

*Purpose: Chloroform removes residual proteins, membrane lipids and detergents by dissolution or accumulation at the aqueous interface. Isoamylalcohol helps produce a clean interface between the aqueous phase containing DNA and the chloroform. If the interface nonetheless falls apart during this step, briefly centrifuge again for 1 min. If the supernatant is clear and free of chloroform or particles after the first chloroform wash, you can omit the second wash and proceed to the next step.*

7. To each DNA extract, add linear polyacrylamide (LPA) to a concentration of 10 µg/mL, and 0.2 volumes of 5M sodium chloride. Mix briefly by inverting 3 times. Then add 2.5 volume ethanol, mix thoroughly, and precipitate in dark at room temperature for 2 hrs.

*Avoid premixing the LPA and ethanol, since it will precipitate the LPA before the LPA has come into contact with DNA.*

8. Centrifuge at room temperature and 14,000×g for 30 min.
9. Decant supernatant, spin the tubes briefly, pipette residual ethanol and let pellets to dry. A small residue of water (moist pellet) poses no problem, and in fact is preferable, since it makes resuspension easier and reduces DNA shearing due to desiccation
10. Resuspend pellet in molecular grade water and let it stand overnight at 4 °C to dissolve.

**Method-3**

(Alternative DNeasy PowerMax Soil Protocol for RNA/DNA from low biomass Soil with low humics)

1. Add 5 grams of soil to the bead tube.

2. Add 5 ml of phenol:chloroform:isoamyl alcohol (25:24:1), pH 8.

3. Add 10 ml PowerBead Solution and 1 ml Solution C1.

4. Homogenize horizontally on a vortex in a 50 ml tube adapter for 10 minutes.

5. Centrifuge 4500 x g for 8 minutes. For centrifuges with low g-force, spin longer. E.g. 2500 x g for 15 minutes.

6. Remove the supernatant. Depending on the soil, there may or may not be a visible upper aqueous layer. If visible, remove only the upper aqueous layer to a new 50 ml tube. It should be about 10 ml. If RNA is not desired add 10 ul of RNase A to the collection tube after the aqueous portion is transferred and incubate at room temperature for 10 minutes.

7. Add 1.5 ml of Solution C2. Cap and shake to mix. Add 1.5 ml of Solution C3. Cap and shake to mix. Centrifuge for 5 minutes at 4500 x g. Note: If the lysate is fairly clear after step 6, then adding 1 ml Solution C2 and 1 ml Solution C3 may be sufficient.

8. Transfer the supernatant to a new tube (~13-14 ml).

9. Add an equal volume (14 ml) of Solution C4. Add 14 ml of 100% ethanol. Mix well (vortex or invert).

10. Load 15 ml of lysate onto the DNeasy PowerMax column.

11. Centrifuge 3 minutes at 4500 x g. Discard the flow-through.

12. Repeat step 10 and 11 until the entire lysate is loaded onto the column.

13. For each prep, prepare the first wash buffer. In a separate tube mix 9 ml of Solution C4 and 11 ml of 100% ethanol. Load onto to the column. Centrifuge 4500 x g for 3 minutes. Discard the flow-through.

14. Wash with 10 ml of Solution C5. Centrifuge 3 minutes at 4500 x g. Discard the flow-through (x2).

15. Wash with 10 ml of 100% ethanol. Centrifuge 3 minutes at 4500 x g. Discard the flow-through (x2).

16. Centrifuge for 10 minutes at 4500 x g in order to dry the column.

17. Place the column into a clean 50 ml tube. Leave the cap off and allow to air dry for 10 minutes further.

18. Elute the DNA/RNA in 6 ml Solution C6 (10 mM Tris-HCl, pH 8.5)

19. Centrifuge 5 minutes at 4500 x g.

20. Optional: Reserve a 50 ul aliquot of eluate and check it on a nanodrop or a gel.

21. To concentrate the DNA/RNA, ethanol precipitate. Add NaCl to a final concentration of 0.2 M. For the expected 5.5 ml eluate, add 240 ul of 5M NaCl. Add 2.5 volumes (14 ml) of 100% ethanol. Optional: add linear acrylamide as a carrier: 10 ul of 5mg/ml. Invert, shake or vortex to mix. Freeze at -20C for at least an hour or overnight.

22. Centrifuge 35 minutes at 4500 x g to pellet the DNA/RNA. Wash the pellet with 70% ethanol (5 ml) and then centrifuge again for 10 minutes at 4500 x g to re-pellet. Decant the ethanol and then turn right side up and allow to air dry until traces of ethanol are mostly gone. Resuspend the pellet in a suitable volume.

**Method-4**

*Remark: Every time the tubes are open ensure that there is no liquid on the lids by applying a short spin*

1. For 5 g of sediment, add 10 mL of Lysis buffer (0.1 M Tris-HCl pH 7.5, 0.05 M EDTA pH 8.0, 1.25 % SDS), 10 µl of RNase (100 mg/ml, Qiagen 19101) and vortex vigorously for 15s (use 50-ml tube).
2. Incubate at 37 °C for 1 hour in a hybridization oven with rotation.
3. Spin samples, add 100 µl Proteinase K (20 mg/ml, ThermoFisher Scientific Cat.No. 25530049) and mix a few times.
4. Incubate at 70 °C for 10 min (statically).
5. Spin samples and add 15 ml of Phenol:Chloform:Isoamyl alcohol (pH 8.05, ThermoFisher Scientific Cat.No. 15593049).
6. Mix thoroughly and centrifuge at 7.500 g at 4 °C for 15 minutes.
7. Transfer aqueous phase into a new 50-ml tube and add 10 ml Chloroform – isoamyl alcohol mixture (24:1).
8. Mix thoroughly and centrifuge at 7.500 g at 4 °C for 10 min.
9. Transfer supernatant to a new 50 ml tube and add 10 µl of LPA (Life Technologies, AM9520) mix well and then add 1/10 volume of 3M sodium acetate (pH 8.0).

10. Add 1 volume of ice-cold Isopropanol and mix thoroughly.
11. Precipitate DNA at -20 °C overnight.
12. Centrifuge at 12.000 g at 4 °C for 35 minutes.
13. Remove supernatant and discard without disturbing the pellet.
14. Wash with 5 ml of 70% Ethanol and centrifuge at 7.500 g at 4 °C for 10-15 minutes.
15. Centrifuge the pellet after washes to collect any residual ethanol.
16. Air-dry the pellet, and elute with 105 ul RNase-free, DNase-free water.
17. Let DNA pellet to dissolve overnight at 4 °C.
